# A Hybrid of Light-Field and Light-Sheet Imaging to Decouple Myocardial Biomechanics from Intracardiac Flow Dynamics

**DOI:** 10.1101/2020.09.08.288480

**Authors:** Zhaoqiang Wang, Yichen Ding, Sandro Satta, Mehrdad Roustaei, Peng Fei, Tzung K. Hsiai

**Author notes:** Correspondence (P.F.); (T.K.H.).

## Abstract

Biomechanical forces intimately contribute to cardiac morphogenesis. However, 4-D (3-D space + time) imaging is needed to investigate the developmental cardiac mechanics with high temporal and spatial resolution. We hereby integrated light-sheet fluorescence microscopy (LSFM) with light-field microscopy (LFM), to simultaneously visualize myocardial contractility and intracardiac blood flow in three dimensions at 200 volumes per second (vps). LSFM allows for reconstruction of the myocardial contraction in zebrafish embryo; and LFM enables simultaneous tracking of the blood cells entering and leaving the contracting heart. We herein established particle tracking velocimetry to interrogate the trajectories of intracardiac blood cells, and we demonstrated deformable image registration to reveal a decrease in the myocardial contractility from atrioventricular (AV) canal to the outflow tract (OFT). We imaged myocardium undergoing torsional contraction and blood flow undergoing regurgitation. Taken together, the integration of light-field and light-sheet microscopy, followed by an image-based analysis pipeline, provides the biomechanical insights into coupling myocardial kinetics with rotational contraction along with intracardiac flow dynamics during development.

## Introduction

Biomechanical forces intimately impart mechanotransduction to modulate cardiac morphogenesis (Bartman et al., 2004; Combs and Yutzey, 2009; Hove et al., 2003; Lee et al., 2013). During development, the myocardium differentiates an outer compact zone and an inner trabeculated zone, and the trabeculated zone is deemed important to facilitate oxygen perfusion and contractile function. (de la Pompa and Epstein, 2012; High and Epstein, 2008; Luxán et al., 2016; Sedmera et al., 2000). We have previously established the multi-scale light-sheet imaging with super-resolution to elucidate the initiation of trabeculation from zebrafish to mouse hearts (Ding et al., 2018; Fei et al., 2019). Using the time-dependent vector analysis and deep-learning approach (Chen et al., 2019), we further enhanced auto-segmentation and quantification of trabecular volume in relation to the compact zone (Ding et al., 2020; Lee et al., 2016; Vedula et al., 2017). Owing to the limited 3-D imaging speed associated with the current optical microscopy, uncoupling myocardial biomechanics from intracardiac flow dynamics to investigate ventricular structure and function remains an imaging challenge.

To this end, we sought to integrate light-field microscopy (LFM) with light-sheet fluorescence microscopy (LSFM) to simultaneously capture the contracting myocardium and intracardiac hemodynamics. The LSFM system was capable of rapidly visualizing the embryonic zebrafish heart at 200 frames per second with high spatial resolution and minimal phototoxicity (Fei et al., 2016; Weber and Huisken, 2015), and an optical gating algorithm synchronized the irregular cardiac periodicity for performing either instantaneous or time-lapse imaging (Lee et al., 2016; Mickoleit et al., 2014; Taylor et al., 2012; Taylor et al., 2019). To track numerous traveling blood cells in space and time, we integrated LFM to image the dynamic signals at single snap shots (Truong et al., 2020; Wagner et al., 2019; Wang et al., 2020). While LSFM acquires a series of plane images across the depth of sample to reconstruct a 3-D image, LFM captures a 3-D image from single 2-D light-field detection. For this reason, light-field requires merely a single exposure for fast volumetric (3-D) detection up to 200 frames per seconds (Prevedel et al., 2014). However, this high temporal resolution is inevitably compromised by a reduced spatial resolution. Thus, the integration of LSFM for high spatial resolution with LFM for fast volumetric detection is complementary to capture cardiac morphogenesis.

In this context, we demonstrate a hybrid imaging system for simultaneous visualization of the contracting cardiomyocytes and traveling blood cells in the beating heart. We developed post-imaging computation to extrapolate the velocity maps, revealing segmental myocardial contraction and intracardiac flow dynamics. We further uncover the distinct magnitudes and directions of contracting myocardium undergoing torsional rotation from AV valves to the outflow track while the traveling blood cells undergoing flow reversal at 2-3 days post fertilization. Integration of light-sheet and light-field microscopy, followed by an automatic post-imaging processing pipeline, further provides a detailed mapping of the developmental cardiac mechanics with high temporal and spatial resolution. Thus, this integration enables the frame-to-frame and dual-channel vector maps to synchronize the interrogation of myocardial kinetics, and tracking of flow trajectories during development.

## Results and Discussion

### Integration of light-field and light-sheet to capture myocardial contraction and intracardiac flow

We demonstrated a sequential imaging pipeline along with a modified retrospective gating approach for dual-channel light-field and light-sheet imaging (**Fig. 1A, Method**). Using the LFM, we tracked the traveling blood cells at up to 200 volumes per second (vps) in the transgenic *Tg(cmlc2:GFP; gata1a:dsRed)* line at 3 dpf. We selectively illuminated the heart by a cylinder-shaped laser beam at a wavelength of 532 nm to eliminate the background noise and to enhance the image contrast (**Fig. 1A: upper panel**) (Truong et al., 2020). Our deep-learning reconstruction algorithm (Wang et al., 2020) allowed for high-throughput reconstruction of the traveling blood cells from the time-dependent light-field sequences. Using LSFM conjugated with a retrospective gating method (Lee et al., 2016; Liebling et al., 2005; Mickoleit et al., 2014; Weber et al., 2017), we performed 3-D reconstruction of the contracting myocardium (**Fig. 1A: lower panel**). The LSFM enabled optical sectioning across the myocardium at 200 fps with high spatial resolution and signal-to-noise ratio (SNR), allowing for detailed analysis of voxel-based displacements (**Fig. 1B**). We reconstructed the 3-D atrium and ventricle, along with the intracardiac blood cells over several cardiac cycles. Using the time-stamps and image-wise similarity function, we synchronized the reconstructed results from both LFM and LSFM at 200 vps to reconstruct a dual-color 4-D beating zebrafish heart (**Figs. 1C, D, Movie 1, 5, 6**). Thus, the dual-channel dataset provides the post-processing pipeline to automatically characterize the myocardial contraction and blood flow.

**Figure 1.**
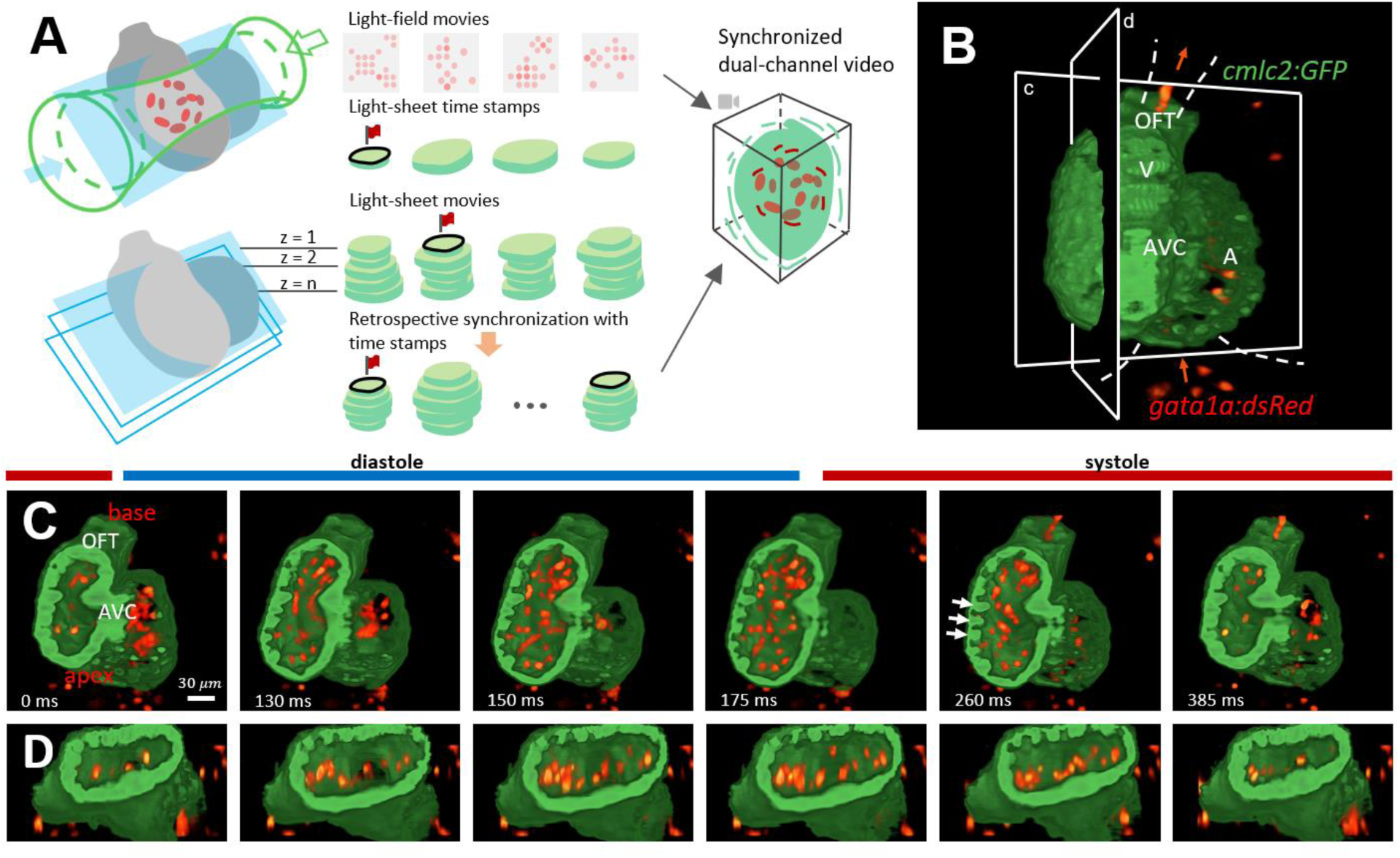
A Pipeline for high speed imaging to couple ventricular contraction with intracardiac flow dynamics at 3 days post pertilization. (A) The integration of light-sheet and light-field microscopy captures the contracting myocardium (*cmlc2:GFP*) and traveling blood cells (*gata1a:dsRed*) at 200 volumes per second. The light field-generated dsRed-labeled cells images (upper illustration) were synchronized with the light sheet-generated cross-sectional myocardial images (lower illustration). The retrospective gating algorithm was used to reconstruct myocardial contraction and relaxation. (B) GFP-labeled cardiomyocyte light chain *(cmlc*) and dsRed-labeled blood cells (*gata1a*) were simultaneously captured in a 3-D reconstructed heart. The red arrows indicate the direction of blood flow. A: atrium; V: ventricle. (C & D) A time-dependent image of cardiac cycle is illustrated in the coronal (C) and sagittal (D) plane, respectively. During diastole (light-blue bar), ventrciular relaxation allowed the ds-Red-labeled cells to enter through the atrial-ventricular (AV) valve. During systole, ventricular contraction drove the cells to leave through the outflow tract (OFT). Morphological changes in endocardial trabeculae (white arrows) are well-delineated during the cardiac cycle.

Despite the sub-optimal spatial resolution for the densely organized cardiomyocytes, LFM demonstrated the capability for tracking the sparsely distributed signals like blood cells (Truong et al., 2020; Wagner et al., 2019; Wang et al., 2020). This is further applicable to study the myocardial Calcium (Ca^2+^) flux via the genetically encoded Ca^2+^ indicators (GECIs) such as GCaMP for electromechanical coupling (Weber et al., 2017) and particle tracers such as nanoparticles (Craig et al., 2012) to extend the application of our method.

### 4-D Vector fields for myocardial contraction

The magnitude and direction of myocardial contractility depend on the biophysical properties, including tissue curvature, sarcomeral length, fiber orientation, myocardial thickness and electrical conductivity (Torrent-Guasp et al., 2005). Images acquired by our pipeline embraced the time- and location-dependent properties for precise vector analyses (**Fig. 2A**). We computed the myocardial displacement between two consecutive frames (**Fig. 2A**) by using deformable image registration (DIR) to infer the voxel-based vector fields (**Fig. 2B**). Each vector indicated the direction and velocity of contractile displacement (μm/s) at corresponding region. A heatmap further reveals spatial variations in the myocardial contractile velocity (**Fig. 2C**). The velocity is elevated toward the apex and AV canal, suggesting an elevated myocardial kinetic energy (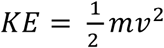, where *m* denotes the mass and *v* the velocity). This accentuated KE in the apex and AV canal further overlaps with the clockwise rotation in Segments 1 and counterclockwise rotation in Segment 2, resulting in shortening of the ventricular long axis during systole (see **Fig. 3D**). This image-based displacement analysis provided a dynamic platform to uncover the spatial and temporal variations in vector fields during myocardial contraction (**Movie 2**).

**Figure 2.**
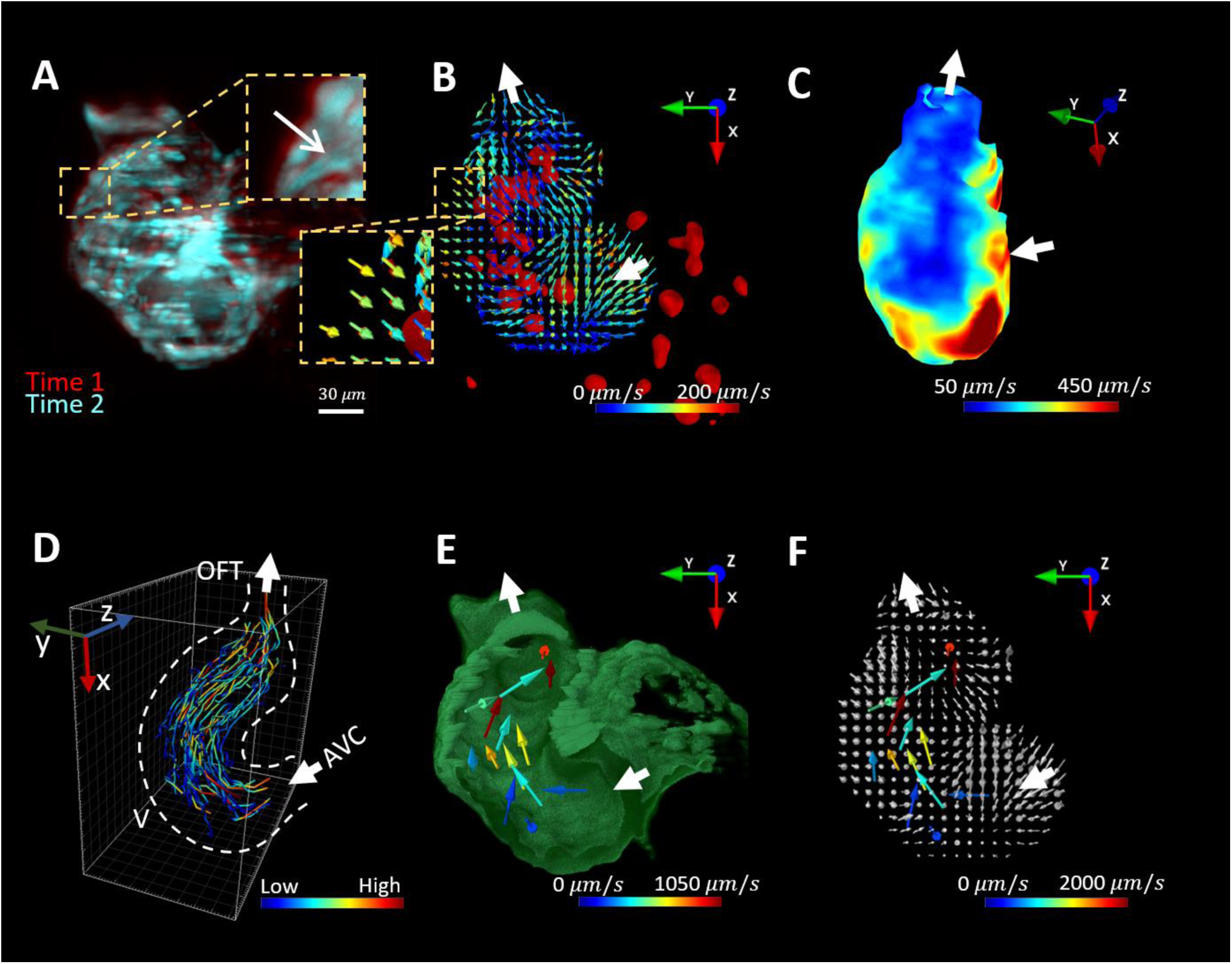
Post-imaging computation to reconstruct 4-D vector fields for myocardial contraction and blood flow. (A) Myocardial contraction from 2 different time points are overlaid to demonstrate myocardial displacement. The arrow indicates the direction of displacement. (B) Deformable image registration was used to infer the vector fields of the myocardial displacement. Each vector indicates the direction and velocity of contraction as color-coded by the magnitude (*μm/s*) in the corresponding voxel. Intracardiac ds-Red labeled cells (red) and direction (white) were superimposed with vector fields. (C) A representative heatmap depecits the magnitudes of contractile velocity of the ventricle. (D) Post-imaging process reveals the trajectories of the individual cells during a cardiac cycle. V: ventricle; OFT: outflow tract; AVC: atrioventricular canal. (E) The vector fields represents the velocity of dsRed-labeled cells. (F) The vector fields for myocardial displacement and the traveling cells were merged to couple myocardial contraction with intracardiac flow dynamics.

**Figure 3.**
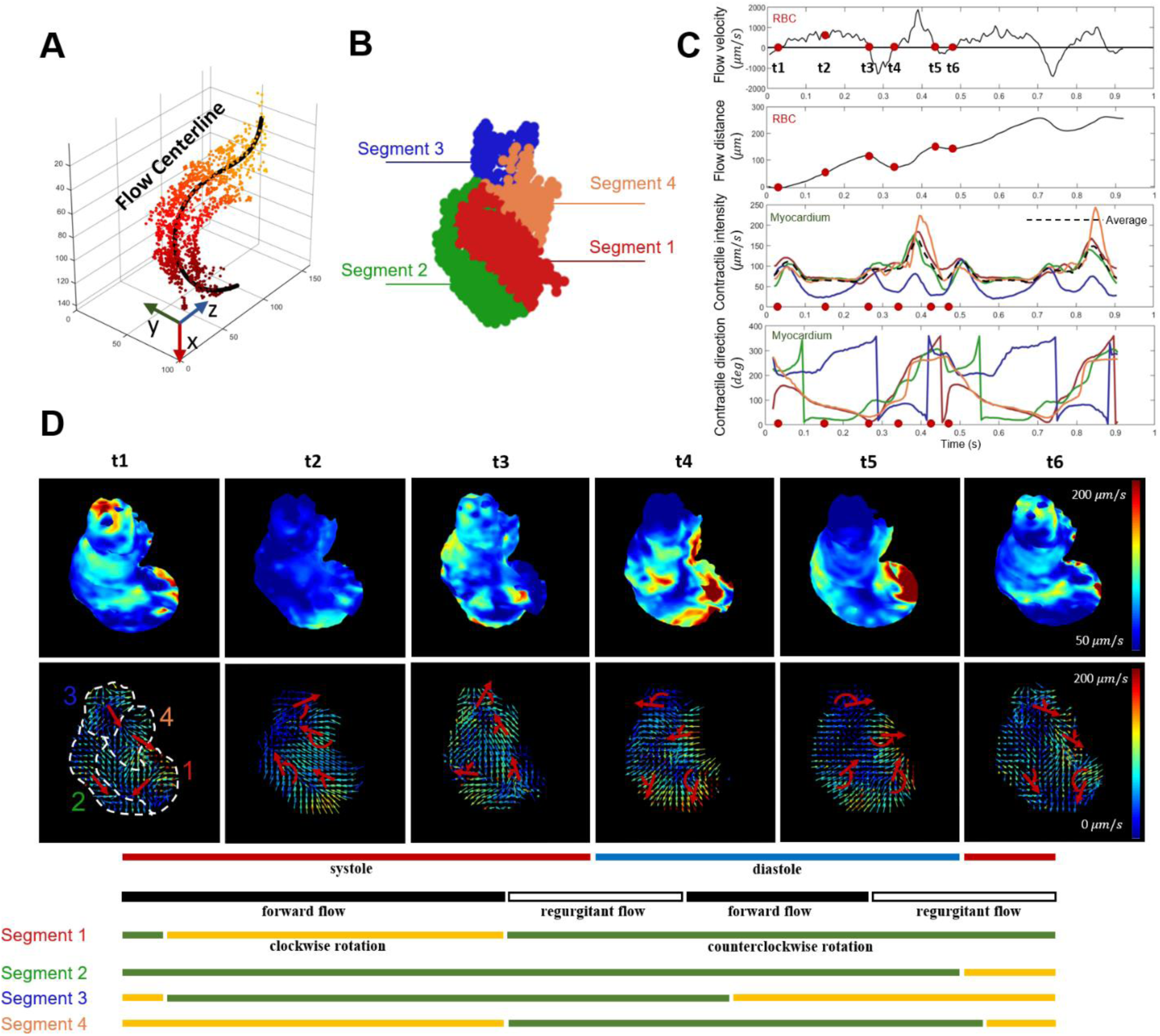
Frame-to-frame analyses of the ventricular contraction and intracardiac flow dynamics at 3 dpf. (A) The mean trajectory of the individual dsRed-labeled blood cells was depicted as the flow centerline, starting from the AV cannel (dark colors) to the outflow tract (light colors). (B) The myocardium was divided into 4 segments to demonstrate variations in vector fields. (C) The velocity of individual blood cells was projected onto the flow centerline, and the mean flow velocity was plotted as a function of cardiac cycles. Myocardial contractility was quantified in terms of the displacement, contraction rate, and direction. Rotational contraction of the myocardium developed in the 4 different segments, ranging from 0 to 360°. (D) Six representative time points (from t1 to t6) reveal the heat maps for the contracting myocardium (upper panels) and vector maps (lower panels) for torsional contraction in the 4 different segments.

We generated the myocardial displacement vector maps by exploiting the gray scale raw images, obviating the need for manual segmentation and annotation. While this enables frame-to-frame analyses of the mechanics with high resolution, endocardiac segmentation would further enhance the accuracy of estimating the volumetric geometry needed to compute metrics such as ventricular ejection fraction (Akerberg et al., 2019; Ding et al., 2020).

### 4-D Vector fields for intracardiac blood flow

To address the intracardiac flow dynamics, we employed the Particle Tracking Velocimetry (PTV) to position the individual cells over two cardiac cycles. We mapped the trajectories of 81 traveling cells from the AV canal to outflow track (OFT) (**Fig. 2D**), and we extracted the velocity vectors of the cells at each frame (**Fig. 2E, Movie 3**). We further demonstrated a heatmap for the 3-D mean velocity distribution of blood flow (**Fig. S4**). Taken together, we integrated DIR with PTV techqniues to merge 1) the vector fields for the myocardial displacement (**Fig. 2B**) with 2) the velocity vectors for the traveling cells (**Fig. 2E**); thereby, establishing the mechanical and hemodynamic coupling (**Fig. 2F**).

Unlike the conventional Particle Image Velocimetry (PIV) (Hove, 2006; Hove et al., 2003; Jamison et al., 2013), PTV has been demonstrated to capture the intracardiac blood flow (Yalcin et al., 2017), allowing for mapping the trajectories of individual dsRed-labeled blood cells from AVC to outflow track (OFT). While PIV provides high spatial resolution and detailed velocity profiles, PTV enables time-dependent tracking of each individual signal in a dynamic space. However, PIV is ineffective in the settings of sparse signals and in narrow space where the size of tracer is comparable to the vessel (Boselli and Vermot, 2016). For these reasons, PTV was chosen in our method. Our results could potentially corroborate the *in silico* simulation of intracardiac flow dynamics and microvascular network (Boselli and Vermot, 2016; Lee et al., 2018; Vedula et al., 2017).

### Frame-to-frame analysis of the 4-D ventricular biomechanics

To quantify the flow pattern, we identified a mean flow direction by fitting a centerline to the distribution of the traveling blood cells (**Fig. 3A and Fig. S3**). We averaged the velocity vectors along the flow centerline to generate a mean velocity (**Fig. 3C: velocity vs. time**). The negative values (>1000 μm/s) at the end of systole and diastole denote flow reversal, resulting in a decrease in the forward flow (**Fig. 3C: distance vs. time**). The flow reversal accounts for 29.8% of systole and 42.5% of diastole duration, and 39.1% and 10.5% for the distance travelled, respectively. These percentages implicate an inefficiency of myocardial kinetics during AV canal formation and outflow track development following cardiac looping at 3dpf.

The ventricle was divided into 4 segments for comparing myocardial contraction and direction in relation to the flow centerline (**Fig. 3B**). We calculated the magnitude-weighted mean vectors to demonstrate the rates of segmental displacement along with the direction (**Fig. 3C: Contraction and Direction vs. time**). As a corollary, we analyzed the spatial and temporal variations in myocardial contraction with the heatmaps that were color-coded with the magnitudes of displacement to demonstrate a gradient of contractile force in various segments (**Fig. 3D, upper panels, Movie 4**). During systole, the contraction rate in the OFT (segment 3) was moderate in comparison to segments 1 (red) and 4 (orange). During diastole, all segments underwent a rapid rate of relaxation. Notably, segments 1 & 4, proximal to the atrium, underwent clockwise rotation during systole and counterclockwise during diastole (**Fig. 3D: lower panels**). We determined the clockwise and counterclockwise rotation by the relative changes rather than the absolute angles. While Segments 2 and 3 underwent counterclockwise contraction (green and blue segments), Segments 1 and 4 underwent clockwise contraction (red and yellow segments). (**Fig. 3D**). Despite these two opposing directions, both Segments 1 (red) and 2 (green) shortened the ventricular long axis during systole and both relaxed in the counterclockwise fashion during diastole. This opposite segmental rotation further engenders a torsional myocardial contraction analogous to the human left ventricular contraction.(McCormick and Tzima, 2016; Omar Alaa Mabrouk et al., 2015) (Young and Cowan, 2012) In addition, this torsional contraction couples with the trajectories of 81 dsRed-labeled cells undergoing forward and reversal flow from the AV cannal to outflow track (see **Fig. 2D**). Thus, our hybrid imaging system, in conjunction with efficient image analyses, allowed for decoupling the myocardial biomechanics and flow dynamics to study the cardiac morphogenesis.

## Materials and methods

### Zebrafish line

The transgenic *Tg(cmlc2:GFP; gata1a:dsRed)* zebrafish (*Danio rerio*) line was used to access the tracers of fluorescence-labeled cardiac myosin light chain and blood cells to decouple the contracting myocardium from blood flow. Zebrafish embryos were harvested from natural mating at the UCLA Zebrafish Core Facility, and they were maintained in the standard E3 medium and supplemented with phenylthiourea (PTU, 0.03%, Sigma Aldrich, MO) after 24 hours post fertilization (hpf) to inhibit melanogenesis. Embryos were anesthetized with tricaine (3-amino benzoic acidethylester, 0.2 mg/mL, Sigma Aldrich, MO), embedded in 1% low-melting-point agarose, and mounted in the Fluorinated Ethylene Propylene (FEP) under the microscope. All animal studies were performed in compliance with IACUC protocol approved by the UCLA Office of Animal Research.

### Integration of light-field and light-sheet imaging

To establish the orthogonal geometry of LSFM, we used a dual illumination system featuring both selective volume and plane illumination for dual-channel detection.(Wang et al., 2020) Two dry objectives (Plan Fluor 4×/0.13, Nikon) for illumination were placed in the opposing direction, and were perpendicularly positioned to the water dipping detection objective (Fluor 20×/0.5w, Nikon). Zebrafish were held by a multi-dimension stage (Newport) to adjust their position and orientation. Zebrafish was oriented and scanned by a stepper motorized actuator (ZST225B, Thorlabs) in the direction of detection axis. For light-field detection, we used a macro lens (AF 60 mm 2.8D, Nikon) to relay the back focal plane of microlens array (MLA, APO-Q-P150-F3.5 (633), OKO Optics) onto the camera sensor. sCMOS cameras (Flash 4.0 V2, Hamamatsu) were installed and synchronized by external trigger (see **Fig. S2**).

### Imaging pipeline and retrospective gating

Light-field and light-sheet imaging were sequentially applied with retrospective gating method in the post-imaging processing for synchronization. Both selective volume and selective plane illumination were simultaneously implemented in our setup. The light-field images of dsRed-labeled cells were recorded by light-field mode while the 2-D cross sectioning of the GFP-labeled myocardium was concurrently captured by the light-sheet mode. A movie is provided to illustrate a time stamp to indicate the cardiac cycles of the acquired light fields. We further imaged the contracting myocardium in each of the 2-D slices throughout the entire 3-D heart by the light-sheet mode. Data were collected at the frame rate of 200 Hz and exposure of 5 ms for both modes. The step size was 2 *μ*m, and 50-70 image sequences were captured to cover the entire heart by the light-sheet imaging. Each image sequence, including light fields, time stamps and light-sheet images, contained 450 frames to cover 4-5 cardiac cycles. We adopted a deep-learning model to provide end-to-end conversion from the 2-D light field measurements to 3-D image stacks, providing a rapid 3-D reconstruction of the dsRed-labeled cells.(Wang et al., 2020) We trained the model with the static confocal images of blood cells in the hearts and vasculatures from 16 zebrafish embryos at 3-4 dpf. Next, we enabled the recovery of a 3-D volume (−50 *μ*m to 50 *μ*m in depth) from the light-field raw data. The final output was registered to the light sheet-acquired images of the contracting myocardium.

We adapted the retrospective gating method to synchronize the contracting myocardium to the identical cardiac cycles.(Lee et al., 2016; Liebling et al., 2005; Mickoleit et al., 2014) Assuming regular periodicity, we sought the proper relative temporal shifts between sequences based on the minimization of the difference function (for which we chose pixel-wise Euclidean distance). Both reconstructed dsRed-labeled cells and myocardium were merged for quantification and visualization.

### Computation of vector mappings

The myocardial contraction was inferred by the intensity-based and non-rigid deformable image registration (DIR). We employed the demons method implemented in MATLAB’s Image Processing Toolbox to estimate an optimal transformation, *T*:(*x, y, z*)→(*x′, y′, z′*), mapping a voxel spanning from the reference field to the moving field. We noted that image registration represents a geometric transformation of the image instead of an intensity transformation. In each voxel, a vector was predicted to represent the motion of the objects in the output transformation map, *T*. The registration was performed on 3 pyramid levels from the coarse to fine image resolution with 50, 25, and 10 iterations. To suppress the errors, we applied an average window of 7 frames (30 ms) to maintain smoothness between the successive maps.

We used an automatic tracking method to map the trajectories of dsRed-labeled cells in Imaris. The 3-D image sequence was resized to an isotropic spatial resolution of 2 *μ*m ×2 *μ*m ×2 *μ*m, followed by a Gaussian blur to remove the artifacts. Region of interest was restricted within the ventricle to decrease the computation cost. Point candidates were detected, and the trajectories were formed by an autoregressive motion algorithm with a max gap size of 1. We filtered the results using a thresholding on track duration, and manual inspection was optional. Vector map indicated that the transient motion of dsRed-labeled cells was computed from the trajectories in MATLAB.

### Quantification of the flow trajectories

To characterize the forward and reversal flow, we fitted a centerline to simplify the inward and outward flow (**Fig. S3**). We collected all the dsRed-labeled cells from the trajectories throughout the entire cardiac cycles. Before fitting a line to these points in 3-D, we performed a Principal Component Analysis (PCA) to find the principal plane in which the points shared the maximum variation. We defined an angle *θ* for each point to a pre-defined center point (red dots in **Fig. S3**) after projection onto this plane. The 3-D coordinates were fitted to this angle variable via the fourth-degree polynomial functions by solving the least square problem. In parallel, a new coordinate system was defined at each point on the flow centerline where the tangent direction was used as the flow direction from AV canal to OFT (**Fig. S3D**). We projected and averaged the vector maps onto the flow direction to obtain a collective flow velocity by frame-to-frame. The “flow distance” (the distance travelled by the dsRed-labeled cells) was computed by the integral of velocity, while the velocity distribution was visualized by collecting the maximum/average magnitudes at the specific positions (**Fig. S4**).

### Analyses of contractile function

The ventricle was divided into segments by using the new coordinate system, as defined by the blood flow centerline (**Fig. S3D**). We analyzed 4 different segments of the contracting myocardium (**Fig. 3B**). In specific, the four segments located on the left/right of the first half/last half of the flow centerline. The contraction was represented by the average vector magnitude, and the contractile direction was defined by the mean vector weighted by their magnitudes in the specific segments. Prior to computation and visualization, we removed the background by binary thresholding of the raw images. The ventricular region was defined by a user-defined distance threshold to the flow centerline.

## Acknowledgements

We thank Yuan (Linda) Dong from UCLA zebrafish facility for her help in fish maintenance.

## Data availability

The code for image post-processing, quantification and visualization is freely available at https://github.com/aaronzq/cardiac. The data is available upon request addressed to (T.K.H.).

## Supplementary materials

**Supplementary Figure 1.**
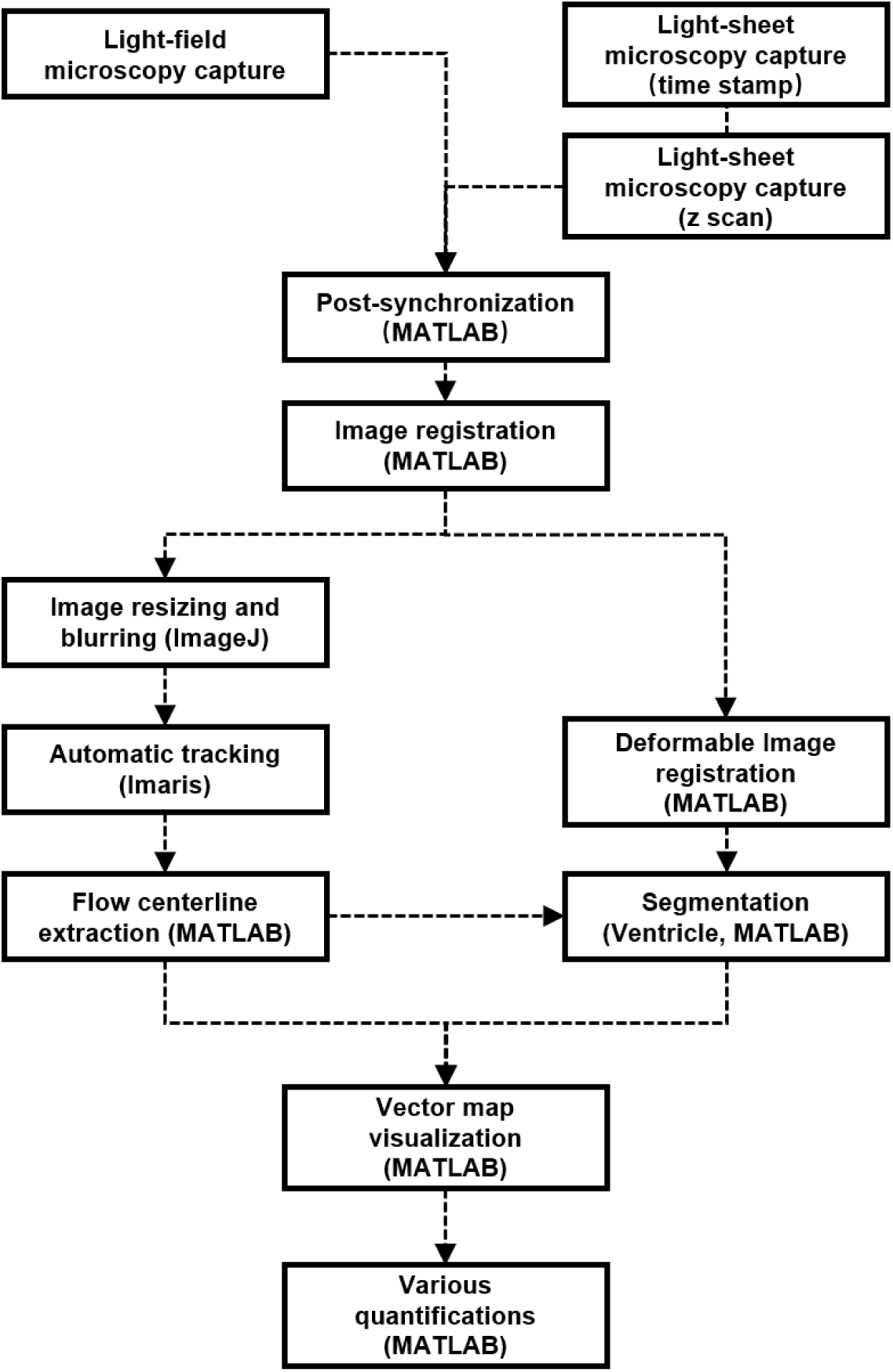
Pipeline for imaging and post-imaging processing. The integration of light-field and light-sheet captures and synchronizes the blood flow and myocardial contraction. Images of dsRed-labeled blood cells have been re-sampled to have 3-D isotropic spatial resolution, followed by Particle Tracking Velocimetry (PTV) to map the flow velocimetry. In parallel, deformable image registration (DIR) allows for displacement analysis of the contracting myocardium. The flow centerline, inferred from blood cells distribution, also indicates the geometry of the ventricle; thereby, facilitating segmentation of the ventricular area for displacement analysis.

**Supplementary Figure 2.**
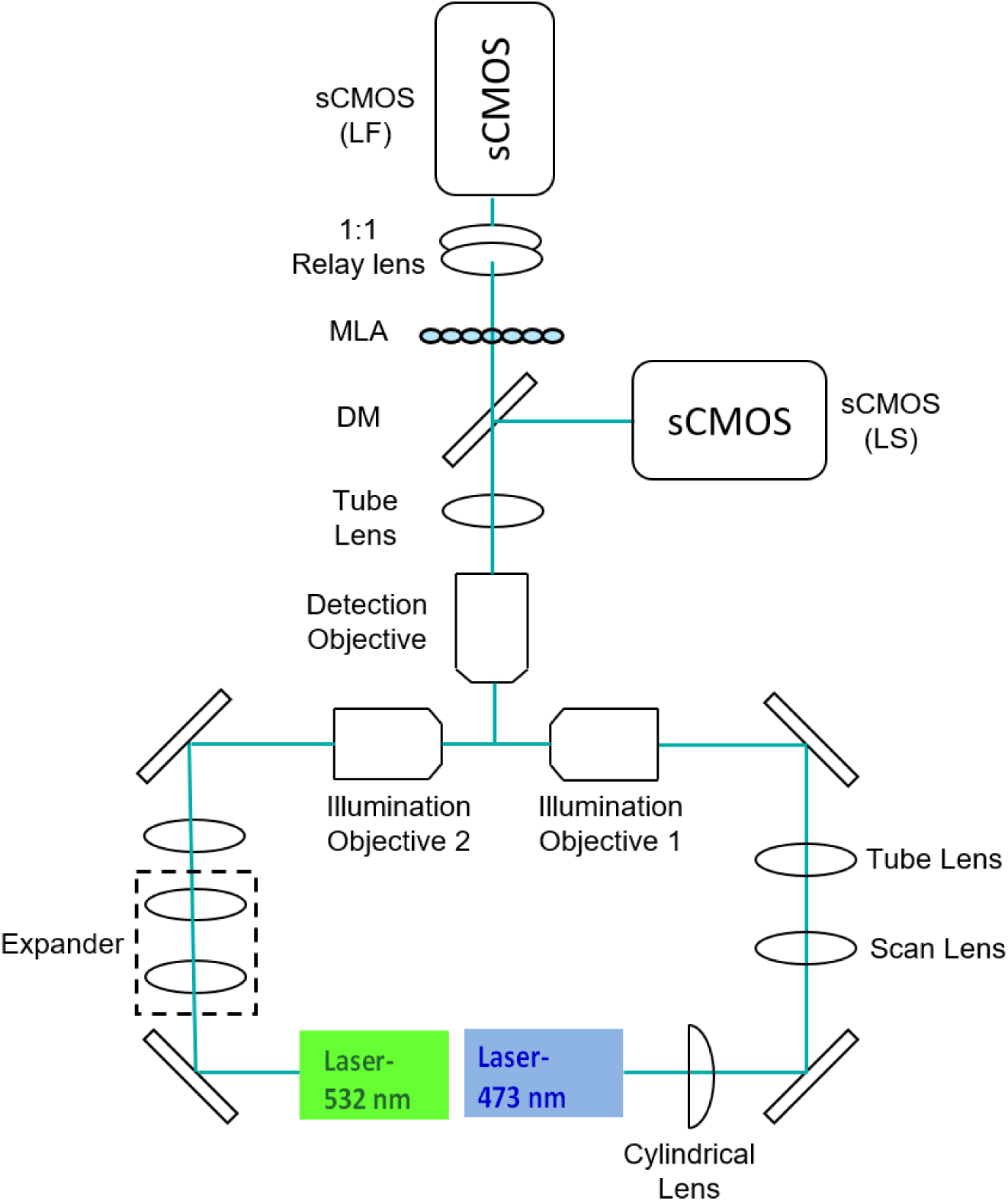
A hybrid setup for dual channel light-field and light-sheet microscopy. Schematic of the integration of light-sheet and light-field system. The laser with different wavelengths (Laser 1 and Laser 2) forms two optical pathways entering the two opposing illumination objectives 1 & 2. Iris, lens pairs and cylindrical lens are used to modulate the dimension and shape of the beam to form 1) the selective volume illumination for light-field microscopy and 2) the selective plane illumination for light-sheet microscopy. The fluorescent signal is collected by the detection lens orthogonal to the illumination. A dichromatic mirror (DM) partitions the detected fluorescent signal onto two detection modalities. For light-field detection, a microlens array (MLA) is placed on the intermediate image plane, and the sCMOS camera is conjugated to the back focal plane of MLA through a 1:1 relay lens. For light-sheet detection, a conventional wide-field is applied.

**Supplementary Figure 3.**
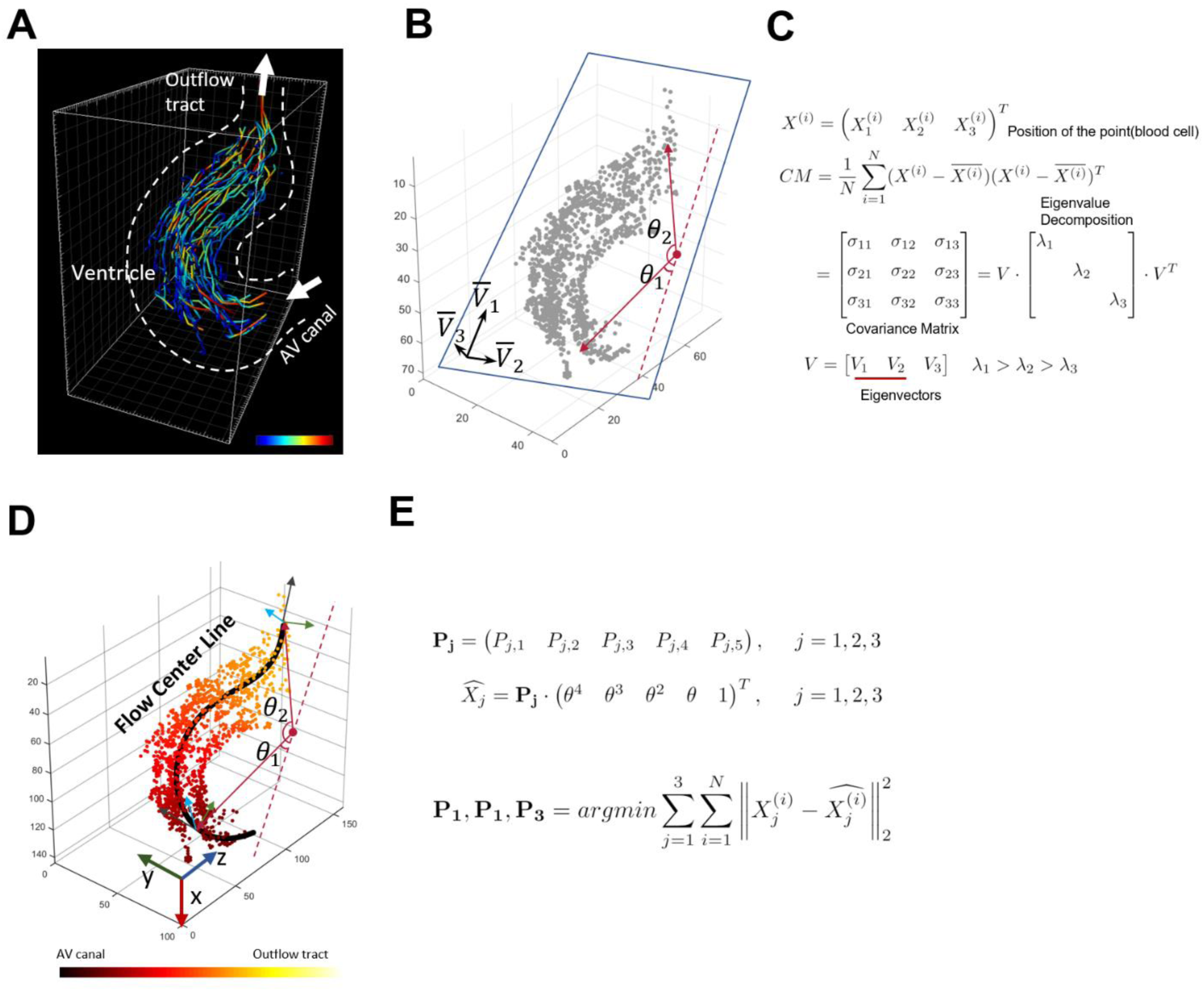
Computation to generate the centerline for ventricular blood flow. (A) Trajectories of the individual blood cells were reconstructed by automatic tracking throughout the cardiac cycle. (B) Principal Component Analysis (PCA) was used to define the main plane where a variation in cell distribution (grey) occurred. The angle of each cell in reference to the center point (red) is defined as *θ*. (C) For the PCA analysis, the covariance matrix was computed on the 3-D coordinates. By eigen value decomposition, we extracted eigen vectors of the first two principal components to define the main plane. (D) A centerline was fitted through the blood cells. For each point along the centerline, we defined a new coordinate system (see axis on the line). (E) Centerline fitting was derived as a least square problem. Each coordinate (x, y or z) of the point *X* is fitted as the fourth order polynomial function with an angle *θ*. Coefficients *P*_1_, *P*_2_, *P*_3_, were computed by solving the minimization problem.

**Supplementary Figure 4.**
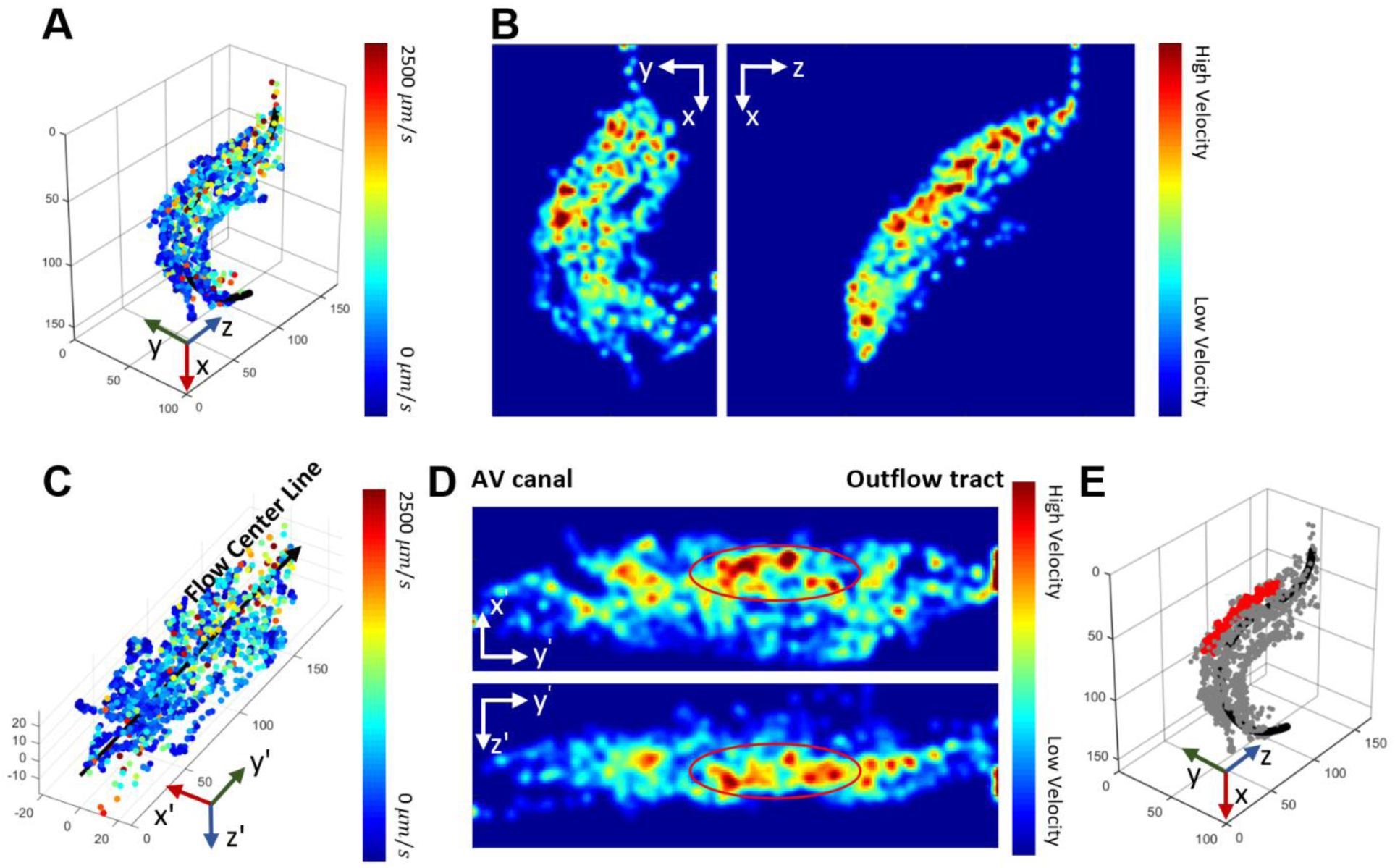
Velocity mapping for the individual dsRed-labeled cells. (A) Mapping of the intracardiac blood cells with color-coded magnitude reveals the distribution of average velocity during a cardiac cycle. (B) Heatmaps were illustrated in the projected views. (C & D) The traveling cells were visualized in the transformed coordinate system with respect to the center flow line. The swapped coordinate system provides a standard perspective to analyze the intracardiac flow dynamics, bypassing the variations from the different imaging orientations. (E) The elevated intracardiac flow velocity was highlighted in red, corresponding to the circled regions in (D).

**Supplementary Figure 5.**
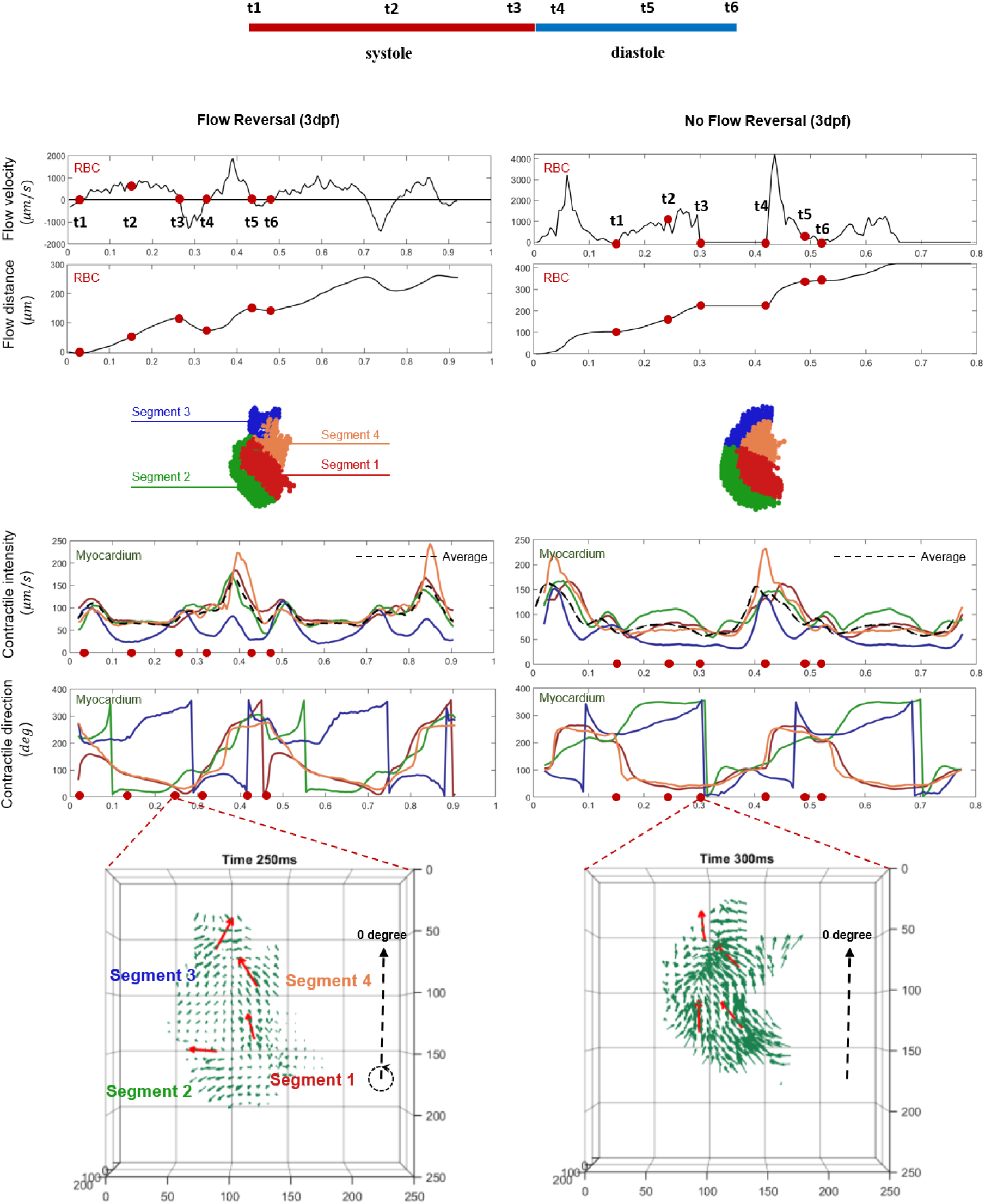
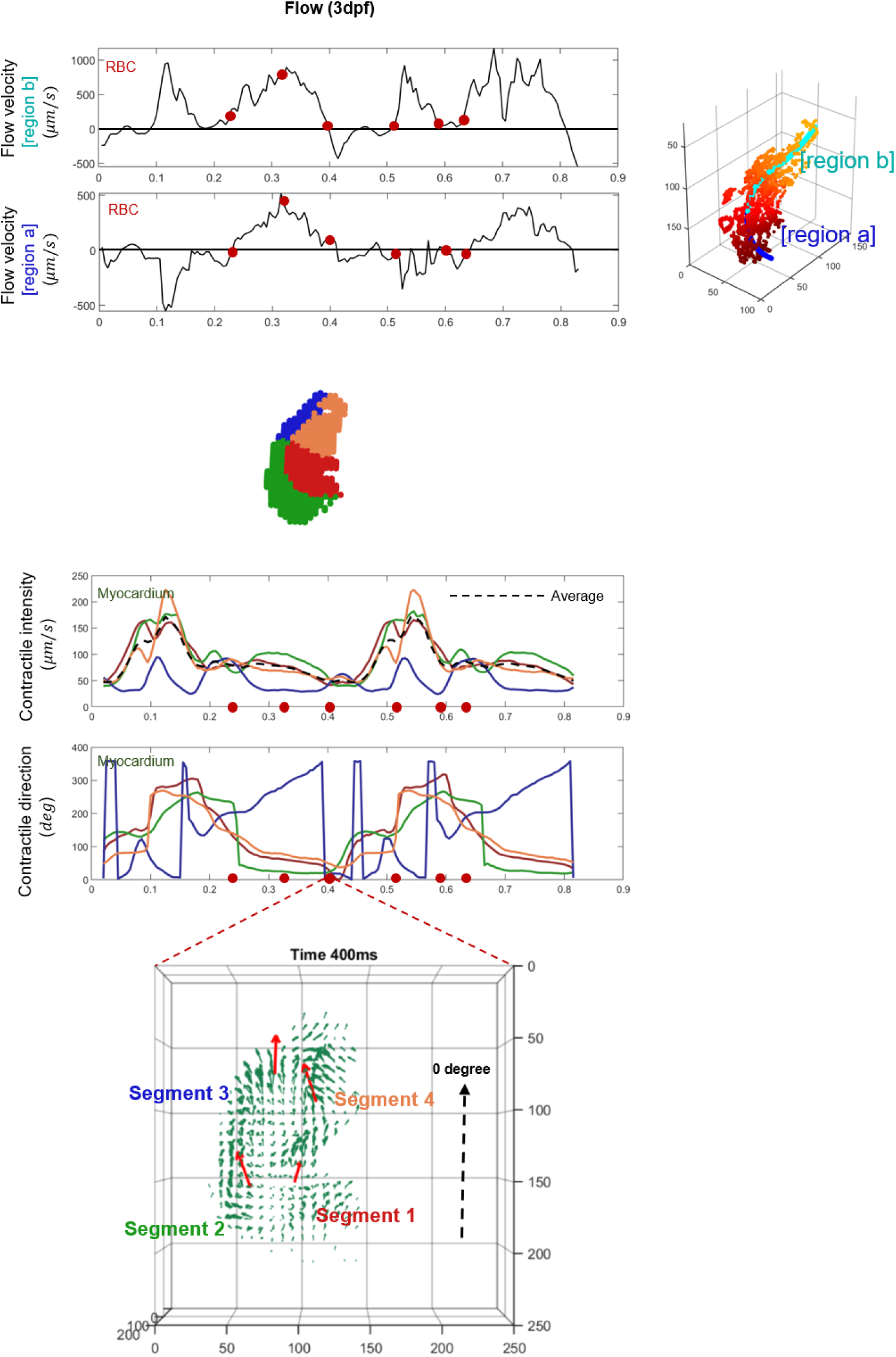
The frame-to-frame analyses of the ventricular contractility and intracardiac flow dynamics from different fish at 3 dpf.

**Supplementary Figure 6.**
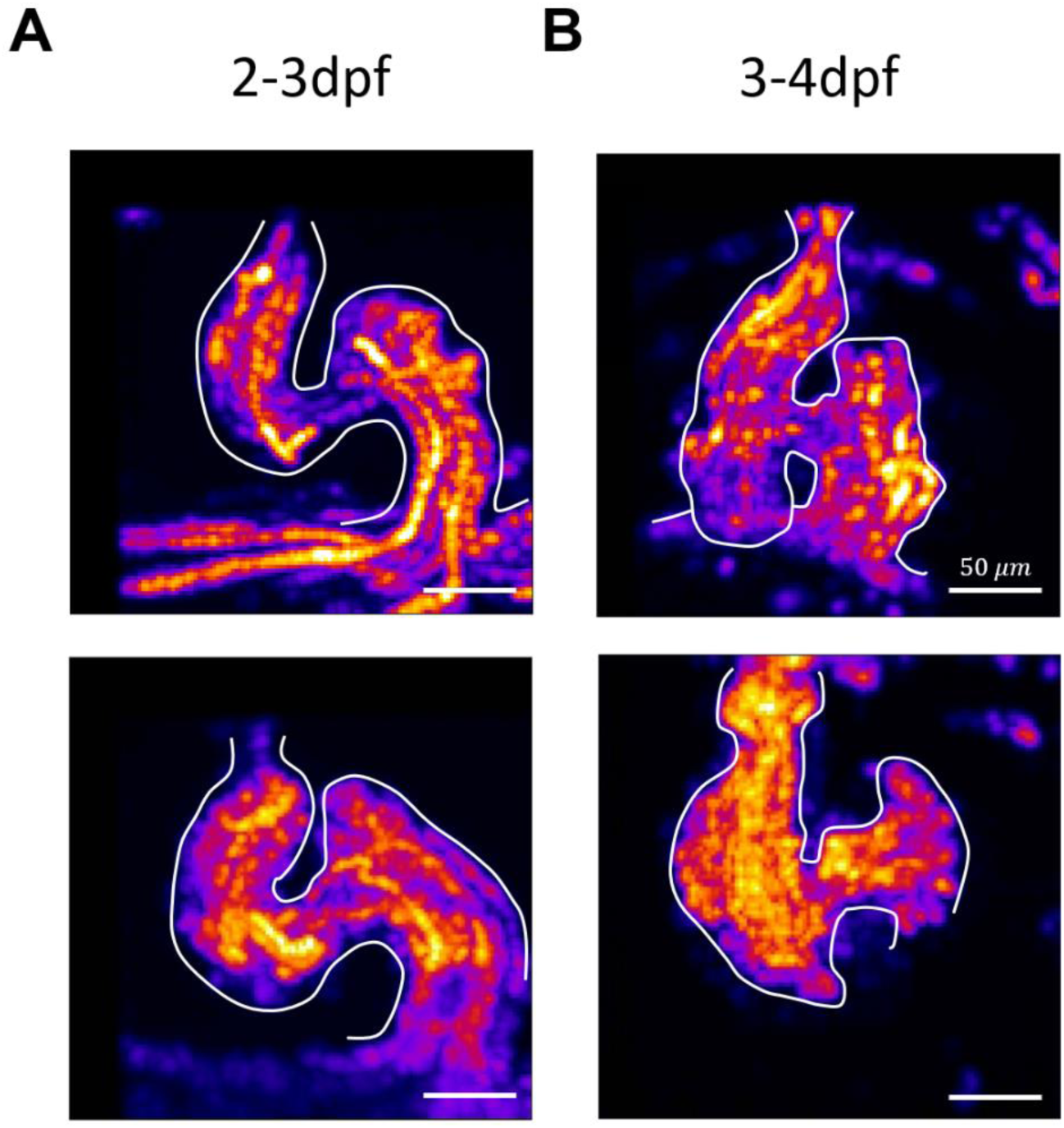
Intracardiac flow domain from 2-3 dpf to 3-4 dpf. Each figure was generated by the summation of different time points over the entire cardiac cycle. (A) Data were acquired during the early stage of cardiac looping (2-3 dpf) and during the (B) late stage (3-4 dpf).

**Movie 1. Dual-color 4-D beating zebrafish heart with flowing blood cells**. Two cardiac cycles have been shown. A, atrium. V, ventricle. Scale bar, 50 *μm*.

**Movie 2. Myocardial displacement during contraction**. Each green vector illustrates the magnitude (vector length) and direction (vector direction) of the displacement of the myocardia in corresponding voxel. Red vectors are mean vectors of the specific segment, after the heart is divided into 4 parts. Grid unit, *μm*.

**Movie 3. Blood cells flowing during contraction**. Blood cells have been tracked and each red vector demonstrates magnitude (vector length) and direction (vector direction) of the motion of each individual tracer. The black line is the flow centerline. Grid unit, *μm*.

**Movie 4. Heatmaps of the myocardial displacement during contraction**. The magnitude of displacement has been encoded in color. Grid unit, *μm*. Color bar unit, *μm/s*.

**Movie 5. Myocardial contraction at different developmental stages at 3 dpf**. Left to right, early to late stage. Scale bar, 50 *μm*.

**Movie 6. Blood flow at different developmental stages at 3 dpf in projection views**. Left to right, early to late stage. Scale bar, 50 *μm*.

## References

Akerberg, A. A., Burns, C. E., Burns, C. G. and Nguyen, C. (2019). Deep learning enables automated volumetric assessments of cardiac function in zebrafish. Dis Model Mech 12, dmm040188.

Bartman, T., Walsh, E. C., Wen, K. K., McKane, M., Ren, J., Alexander, J., Rubenstein, P. A. and Stainier, D. Y. (2004). Early myocardial function affects endocardial cushion development in zebrafish. PLoS Biol 2, E129.

Boselli, F. and Vermot, J. (2016). Live imaging and modeling for shear stress quantification in the embryonic zebrafish heart. Methods (San Diego, Calif.) 94, 129–134.

Chen, J., Ding, Y., Chen, M., Gau, J., Jen, N., Nahal, C., Tu, S., Chen, C., Zhou, S., Chang, C. C., et al. (2019). Displacement analysis of myocardial mechanical deformation (DIAMOND) reveals segmental susceptibility to doxorubicin-induced injury and regeneration. JCI Insight 4.

Combs, M. D. and Yutzey, K. E. (2009). Heart valve development: regulatory networks in development and disease. Circulation research 105, 408–421.

Craig, M. P., Gilday, S. D., Dabiri, D. and Hove, J. R. (2012). An optimized method for delivering flow tracer particles to intravital fluid environments in the developing zebrafish. Zebrafish 9, 108–119.

de la Pompa, J. L. and Epstein, J. A. (2012). Coordinating tissue interactions: Notch signaling in cardiac development and disease. Developmental cell 22, 244–254.

Ding, Y., Gudapati, V., Lin, R., Fei, Y., Packard, R. R. S., Song, S., Chang, C. C., Baek, K. I., Wang, Z., Roustaei, M., et al. (2020). Saak Transform-Based Machine Learning for Light-Sheet Imaging of Cardiac Trabeculation. IEEE transactions on bio-medical engineering.

Ding, Y., Ma, J., Langenbacher, A. D., Baek, K. I., Lee, J., Chang, C. C., Hsu, J. J., Kulkarni, R. P., Belperio, J., Shi, W., et al. (2018). Multiscale light-sheet for rapid imaging of cardiopulmonary system. JCI Insight 3.

Fei, P., Lee, J., Packard, R. R., Sereti, K. I., Xu, H., Ma, J., Ding, Y., Kang, H., Chen, H., Sung, K., et al. (2016). Cardiac Light-Sheet Fluorescent Microscopy for Multi-Scale and Rapid Imaging of Architecture and Function. Sci Rep 6, 22489.

Fei, P., Nie, J., Lee, J., Ding, Y., Li, S., Zhang, H., Hagiwara, M., Yu, T., Segura, T., Ho, C.-M., et al. (2019). Subvoxel light-sheet microscopy for high-resolution high-throughput volumetric imaging of large biomedical specimens. Advanced Photonics, 016002.

High, F. A. and Epstein, J. A. (2008). The multifaceted role of Notch in cardiac development and disease. Nature reviews. Genetics 9, 49–61.

Hove, J. R. (2006). Quantifying cardiovascular flow dynamics during early development. Pediatric research 60, 6–13.

Hove, J. R., Köster, R. W., Forouhar, A. S., Acevedo-Bolton, G., Fraser, S. E. and Gharib, M. J. N. (2003). Intracardiac fluid forces are an essential epigenetic factor for embryonic cardiogenesis. Nature 421, 172–177.

Jamison, R. A., Samarage, C. R., Bryson-Richardson, R. J. and Fouras, A. (2013). In vivo wall shear measurements within the developing zebrafish heart. PLoS One 8, e75722.

Lee, J., Fei, P., Packard, R. R. S., Kang, H., Xu, H., Baek, K. I., Jen, N., Chen, J., Yen, H., Kuo, C. C. J., et al. (2016). 4-Dimensional light-sheet microscopy to elucidate shear stress modulation of cardiac trabeculation. The Journal of Clinical Investigation 126, 1679–1690.

Lee, J., Moghadam, M. E., Kung, E., Cao, H., Beebe, T., Miller, Y., Roman, B. L., Lien, C.-L., Chi, N. C., Marsden, A. L., et al. (2013). Moving domain computational fluid dynamics to interface with an embryonic model of cardiac morphogenesis. PLoS One 8, e72924–e72924.

Lee, J., Vedula, V., Baek, K. I., Chen, J., Hsu, J. J., Ding, Y., Chang, C. C., Kang, H., Small, A., Fei, P., et al. (2018). Spatial and temporal variations in hemodynamic forces initiate cardiac trabeculation. JCI Insight 3.

Liebling, M., Forouhar, A. S., Gharib, M., Fraser, S. E. and Dickinson, M. E. (2005). Four-dimensional cardiac imaging in living embryos via postacquisition synchronization of nongated slice sequences. Journal of biomedical optics 10, 054001.

Luxán, G., D’Amato, G., MacGrogan, D. and de la Pompa, J. L. (2016). Endocardial Notch Signaling in Cardiac Development and Disease. Circulation research 118, e1–e18.

McCormick, M. E. and Tzima, E. (2016). Pulling on my heartstrings: mechanotransduction in cardiac development and function. Current opinion in hematology 23, 235–242.

Mickoleit, M., Schmid, B., Weber, M., Fahrbach, F. O., Hombach, S., Reischauer, S. and Huisken, J. (2014). High-resolution reconstruction of the beating zebrafish heart. Nat Methods 11, 919–922.

Omar Alaa Mabrouk, S., Vallabhajosyula, S. and Sengupta Partho, P. (2015). Left Ventricular Twist and Torsion. Circulation: Cardiovascular Imaging 8, e003029.

Prevedel, R., Yoon, Y. G., Hoffmann, M., Pak, N., Wetzstein, G., Kato, S., Schrödel, T., Raskar, R., Zimmer, M., Boyden, E. S., et al. (2014). Simultaneous whole-animal 3D imaging of neuronal activity using light-field microscopy. Nat Methods 11, 727–730.

Sedmera, D., Pexieder, T., Vuillemin, M., Thompson, R. P. and Anderson, R. H. (2000). Developmental patterning of the myocardium. The Anatomical record 258, 319–337.

Taylor, J. M., Girkin, J. M. and Love, G. D. (2012). High-resolution 3D optical microscopy inside the beating zebrafish heart using prospective optical gating. Biomed Opt Express 3, 3043–3053.

Taylor, J. M., Nelson, C. J., Bruton, F. A., Kaveh, A., Buckley, C., Tucker, C. S., Rossi, A. G., Mullins, J. J. and Denvir, M. A. (2019). Adaptive prospective optical gating enables day-long 3D time-lapse imaging of the beating embryonic zebrafish heart. Nat Commun 10, 5173.

Torrent-Guasp, F., Kocica, M. J., Corno, A. F., Komeda, M., Carreras-Costa, F., Flotats, A., Cosin-Aguillar, J. and Wen, H. (2005). Towards new understanding of the heart structure and function. European journal of cardio-thoracic surgery : official journal of the European Association for Cardio-thoracic Surgery 27, 191–201.

Truong, T. V., Holland, D. B., Madaan, S., Andreev, A., Keomanee-Dizon, K., Troll, J. V., Koo, D. E. S., McFall-Ngai, M. J. and Fraser, S. E. (2020). High-contrast, synchronous volumetric imaging with selective volume illumination microscopy. Communications Biology 3, 74.

Vedula, V., Lee, J., Xu, H., Kuo, C. J., Hsiai, T. K. and Marsden, A. L. (2017). A method to quantify mechanobiologic forces during zebrafish cardiac development using 4-D light sheet imaging and computational modeling. PLoS computational biology 13, e1005828.

Wagner, N., Norlin, N., Gierten, J., de Medeiros, G., Balázs, B., Wittbrodt, J., Hufnagel, L. and Prevedel, R. (2019). Instantaneous isotropic volumetric imaging of fast biological processes. Nat Methods 16, 497–500.

Wang, Z., Zhang, H., Zhu, L., Li, G., Li, Y., Yang, Y., Ding, Y., Zhen, M., Gao, S., Hsiai, T. K., et al. (2020). Network-based instantaneous recording and video-rate reconstruction of 4D biological dynamics. bioRxiv, 432807.

Weber, M. and Huisken, J. (2015). In vivo imaging of cardiac development and function in zebrafish using light sheet microscopy. Swiss medical weekly 145.

Weber, M., Scherf, N., Meyer, A. M., Panáková, D., Kohl, P. and Huisken, J. (2017). Cell-accurate optical mapping across the entire developing heart. eLife 6, e28307.

Yalcin, H. C., Amindari, A., Butcher, J. T., Althani, A. and Yacoub, M. (2017). Heart function and hemodynamic analysis for zebrafish embryos. Developmental dynamics : an official publication of the American Association of Anatomists 246, 868–880.

Young, A. A. and Cowan, B. R. (2012). Evaluation of left ventricular torsion by cardiovascular magnetic resonance. Journal of cardiovascular magnetic resonance : official journal of the Society for Cardiovascular Magnetic Resonance 14, 49.

